# Estimating the evolutionary fitness of specific synonymous codon changes

**DOI:** 10.64898/2026.03.16.712166

**Authors:** Vitor A. C. Pavinato, Jody Hey

## Abstract

Synonymous mutations do not alter proteins but undergo natural selection in many species. For *Drosophila melanogaster*, reports on selection strength vary from undetectable to surprisingly strong. Here we apply a new method to estimate the population selection coefficient (2 *Ns*) for all 134 ordered pairs of synonymous codon changes. The method uses ratios of site frequency spectra (SFS) for codon changes to neutral changes, and does not depend on divergence data, or codon frequencies, and is relatively insensitive to demographic history. Results indicate that natural selection on synonymous codons is weak, with |2 *Ns*|<2.07 for all pairs of codons and |2 *Ns*|<1 for 64% of codon changes. Despite being derived solely from polymorphism data, codon fitness estimates are strongly correlated with observed codon frequencies. A selection-mutation-drift model based on our 2 *Ns* estimates accurately predicts codon usage, while a model based on mutation alone fails. Codon fitnesses correlate strongly with a measure of codon frequency covariation among genes, and codons with large frequency differences between high- and low-expression genes have high fitness values. Finally, we detect clear signs that selection favors codon changes that stabilize mRNA secondary structure. By avoiding divergence data, which have multiple sources of additional variance, and focusing solely on allele frequencies, a clear picture emerges of selection on codon usage. The convergence of multiple independent lines of evidence (codon frequencies, expression-dependent usage, covariation patterns, and mRNA structure) validates this polymorphism-based approach and provides a coherent framework for understanding selection on synonymous site evolution.

**Significance statement:** Synonymous mutations do not change proteins yet affect gene expression in multiple ways. Here we provide the first fitness estimates for each type of synonymous codon change. Using only allele frequencies—without divergence data, codon counts, or assumptions about preferred codons—we estimate the fitness of all 134 single-step synonymous codon changes in *Drosophila melanogaster*. Results reveal weak but non-negligible selection, with codon fitness values accurately predicting codon usage patterns, including expression-dependent biases, codon covariation across genes, and mRNA secondary structure effects. This work offers the first comprehensive fitness estimates for individual synonymous mutations, providing a direct population-level view of how subtle genetic changes shape molecular evolution and establishing a general strategy for quantifying weak selection using polymorphism data alone.

## Introduction

Although they do not alter proteins, synonymous mutations are not selectively neutral in many species. The evidence for this comes from multiple sources including experimental evolution studies (Bailey, et al. 2021), a wide array of correlations with diverse aspects of gene expression (Plotkin and Kudla 2011), phylogenetic studies (McVean and Vieira 2001; Singh, et al. 2007) and population genetic analysis (Lawrie, et al. 2013). As expected from theory, the evidence tends to be strongest in species that are thought or known to have large effective population sizes (Hershberg and Petrov 2008).

Synonymous mutations offer a unique opportunity to study natural selection because some aspects of their impact on gene expression are partly understood, and because they provide a form of replication that other classes of variants do not offer. Synonymous mutations of a given codon-to-codon class (e.g., CCC→CCG) are expected to influence gene expression in similar ways through a shared set of molecular mechanisms affecting gene expression (Liu, et al. 2021). This commonality is only approximate, as local effects can arise from gene regulatory differences, mRNA secondary structure, variation in local tRNA pools, and other contextual factors. However, there are no comparably crisp systems for classifying non-synonymous or regulatory mutations by shared mechanisms or expected effect.

We take advantage of the commonality among classes of synonymous variants by applying a new method for estimating the strength of selection for each of the 134 pairs of ancestral/descendant synonymous codon pairs that are one mutational step apart. Figure 1A shows an example of the cumulative site-frequency spectrum for one ordered synonymous SNPs pair for phenylalanine from the autosomes of aligned haploid genomes of a *Drosophila melanogaster* population from Zambia. Shown with each spectrum is a partner neutral spectrum from short intron single nucleotide polymorphisms (SNPs) of the same mutational class (i.e., C to T or T to C) and same flanking base sequences. Also shown is the spectrum for non-synonymous SNPs, to provide perspective. SNPs with deleterious derived alleles will have fewer high frequency alleles (i.e., a cumulative curve above the neutral control), whereas SNPs with beneficial derived alleles will have more high frequency alleles and a cumulative curve below the neutral control. In the case of both ordered changes for phenylalanine codons, Figure 1A shows how the synonymous changes in one direction (TTC→TTT) are above the partner neutral control (i.e. consistent with negative selection) and changes in the other direction (TTT→TTC) are below the neutral control (i.e. consistent with positive selection).

**Figure 1.**
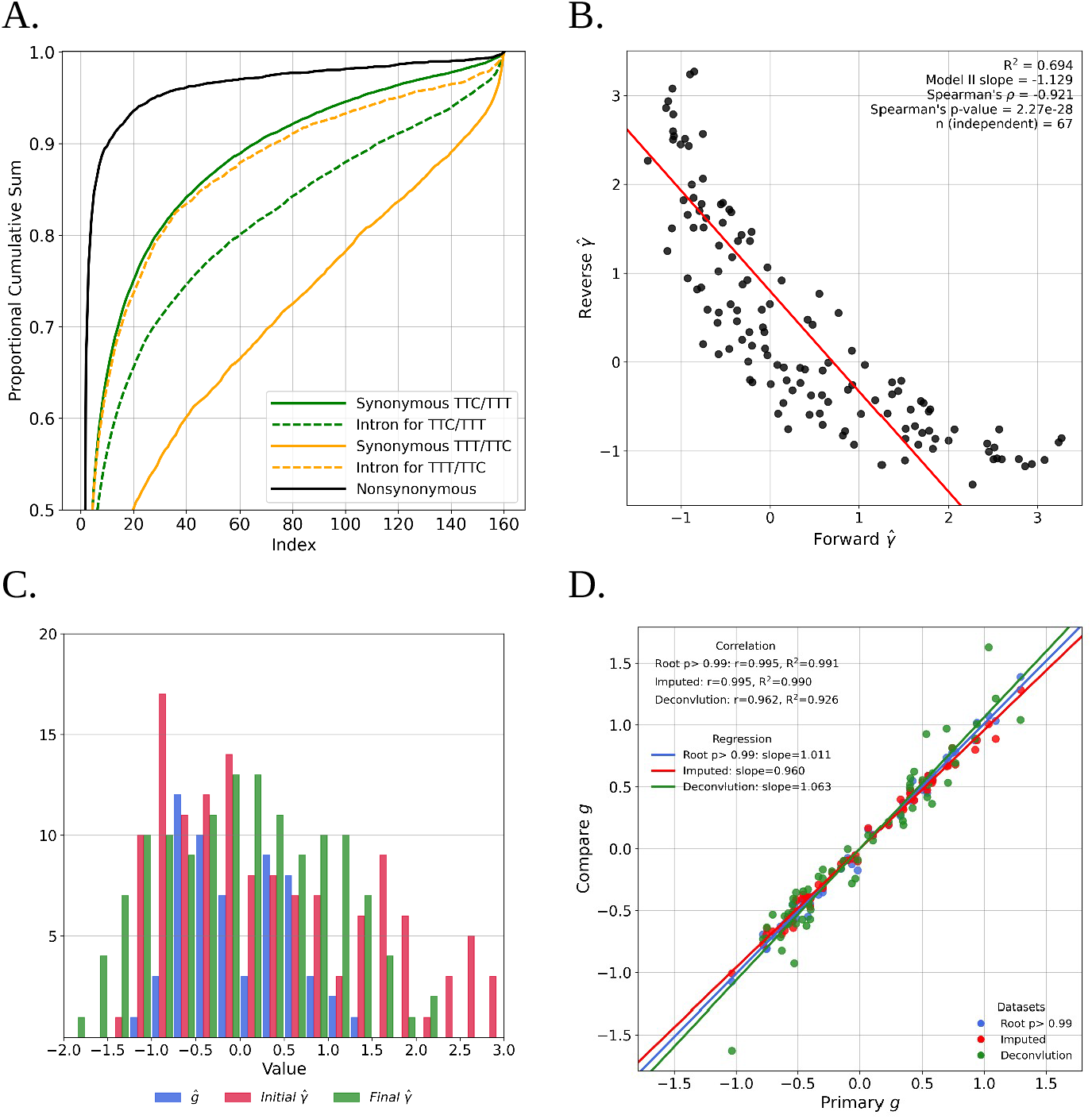
**A.** Cumulative site frequency spectra (as a proportion of the total) for both directions of an ordered codon pair together with paired short intron data. Also shown is the spectrum for non-synonymous SNPs. **B**. Correlation between estimates of forward and reverse codon changes 2 *Ns* values. **C**. Histograms of *g*, initial and final 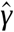 values. **D**. Comparison of alternative pipelines for generating final 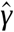 values.

Because both spectra for each ordered codon pair - synonymous and matched intron SNPs - will be affected similarly by demographic and other non-selective factors (e.g. mutation and gene conversion), particularly if selection is weak, 2 *Ns* for the synonymous SNPs can be estimated with a relatively simple likelihood of the ratios of selected (synonymous) to neutral (intron) sites. Our method based on these likelihoods also has the advantage of not relying upon divergence data or codon frequencies (Hey and Pavinato 2025).

## Results

We used a panel of 197 haploid *D. melanogaster* genomes from Zambia (Lack, et al. 2015). Variants were rooted using maximum-likelihood estimates of the *D. melanogaster* ancestral base from an alignment of *melanogaster* group genomes. Because of some missing data, random down-sampling to a sample size of 160 was used to generate the unfolded site frequency spectrum for each of the 134 ordered pairs of codons as well as a companion site frequency spectrum (SFS) from short introns for each ordered codon pair We then applied the SFRatios method to the ratios of SFS counts for codon changes over intron changes (Hey and Pavinato 2025). This approach depends only on the ratios of frequencies of segregating variants and does not depend on any measures of divergence between species, nor any measure of codon usage or codon bias.

For two alleles under weak selection with additive fitness, allele frequency dynamics are a function of the population selection coefficient, 2 *Ns* (denoted hereafter as *γ*), where *N* is the effective population size and *s* is the selection coefficient. Under weak selection, the sign of 2 *Ns* depends on whether the ancestral allele is favored or disfavored relative to the derived allele (Bulmer 1991). Thus each synonymous codon pair can provide two estimates of *γ*, with one ordered SFS providing a negative value and the reverse ordered SFS a positive value (Supplementary Information Table 3).

Because both the forward and reverse values are estimated quantities with measurement uncertainty, a conventional linear regression is not ideal to evaluate if the reciprocity of codon pairs selection strength, since it assumes the predictor is measured without error. Instead, we used a Model II regression with equal error variances, which treats error in both axes symmetrically and is therefore appropriate for testing the expected reciprocal relationship. The full set of 134 reciprocal pairs are not fully independent as each estimate of *γ* appears twice, requiring that the regression was based on an *n* of 67. The fitted model was *Reverseγ* =*α* + *βForwardγ*, with parameter estimates *β*=− 1.129 and 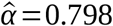. The slope was not significantly different from the theoretical value *β*=− 1 (p = 0.195), and bootstrap inference (200,000 resamples of independent pairs) gave a 95% confidence interval for the slope of [−1.346, −0.934]. In contrast, the intercepts (X: 0.707, Y: 0.798) were significantly greater than zero on both axes (*X* : *p* ≈ 1.0 × 10^−5^, *Y* : *p* ≈ 1.0 × 10^−5^). Our interpretation is that the data are consistent with the general form of the model, i.e., an approximately inverse linear relationship with slope near (−1), but with a significant nonzero offset reflecting estimator bias (Figure 1B).

To obtain internally consistent and less biased *γ* estimates, we constructed a least-squares estimator of a measure of codon fitness, *g*, for each of the 59 codons that can have synonymous mutations. The method assumes only that (1) the forward *γ* is the additive inverse of the reverse *γ*, and (2) the mean fitness change across all codon changes for a given amino acid is zero. Final estimates of *γ* were then generated by taking the difference in *ĝ* values for ordered pairs of codons. The distributions of *ĝ* and initial and final 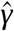 values are shown in Figure 1C (see initial and fitted 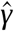 values and *ĝ* values in on Supplementary Information Tables 3, 4 and 5). Values of final 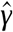 range from −2.07 to 2.07 with mean 95% confidence interval of 0.69, with 86 of the 134 ordered codon pairs having final 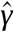 value between −1 and 1, the interval of values often considered to be *effectively* neutral.

Several variations on the *g* estimation pipeline were evaluated, including: (1) using a rooting probability of 0.99 (in contrast to 0.9) when estimating ancestral bases; (2) using a full sample size of 197 genomes without down-sampling, with missing data imputed using Beagle (Browning and Browning 2007); and (3) using data simulated from the actual data to mimic a 5% ancestral polarization error rate using a deconvolution algorithm. All three of these variations generated *ĝ* values very similar to the original values (Supplementary Information Table 6) (Figure 1D).

We then examined the correlation of *ĝ* with codon usage, measured as the frequency of a codon found in protein coding sequences, centered by subtracting the mean frequency across synonymous codons for the same amino acid. The normalization is required to put all codons on the same scale, regardless of the redundancy level of their respective amino acid. Although *ĝ* is based on only relative frequencies of synonymous variants, it is strongly correlated with codon frequency (Figure 2A).

**Figure 2.**
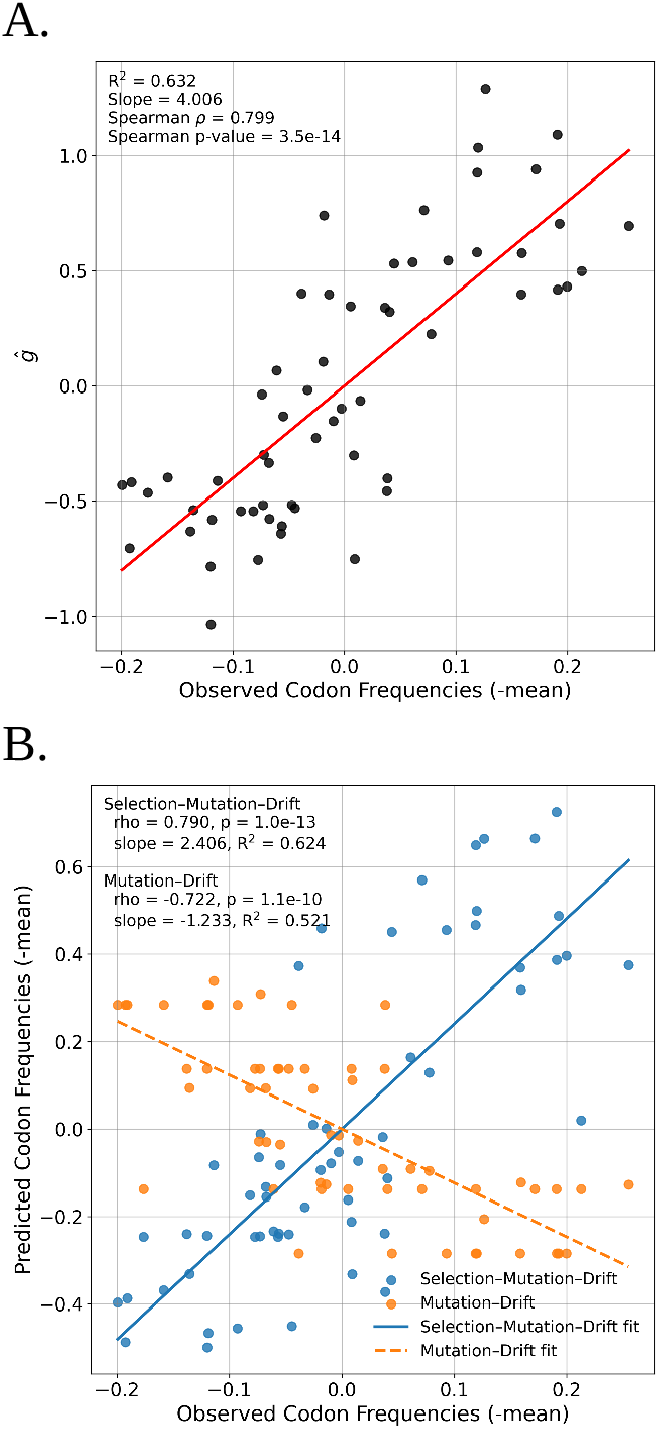
**A.** Comparison between codon fitness (*ĝ*) and the normalized observed codon frequency in the *D. melanogaster* reference genome. **B**. Association between the observed and the expected normalized codon frequencies under a model with selection and mutation and a model with only mutation.

To further examine the associations with codon frequency we predicted codon frequencies under both a selection-mutation-drift model (based on fitted 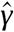 values as well as estimated relative mutation rates) and a mutation-drift model that includes only the relative mutation rates. The model with selection predicted actual codon frequencies far better than the model with only mutation (Figure 2B). In fact, there is a negative association between the predicted and observed values for the model without selection. This is not unexpected because of the known mismatch between the base values of codons that occur at high frequency and the known mutation spectrum in *D*. melanogaster. Most of the more common codons end in G or C (Vicario, et al. 2007), whereas the estimated base-pair mutation matrix has higher rates from GC pairs to AT pairs rather than the reverse (Assaf, et al. 2017). If mutation alone were the primary driver of synonymous codon frequencies then we would expect common codons to typically end in A or T.

As in many species, the most common phenotype that has been associated with variation in codon usage in *D. melanogaster* is gene expression, with selection apparently acting more strongly in genes that are highly expressed (Sharp and Li 1986; Duret and Muchiri 1999). But gene expression is a crude measure that, while correlated with various measures of codon usage in *D. melanogaster*, may obscure important effects in individual tissues (Payne and Alvarez-Ponce 2019; Allen, et al. 2022) or life stages (Vicario, et al. 2008). Codon usage in *D. melanogaster* has also been shown to have effects at many steps between transcription and protein, including translation speed (Cameron, et al. 1999; Zhao, et al. 2017; Wu, et al. 2024), translational accuracy (Akashi 1994; Wu, et al. 2024), modulation of co-translational folding of proteins (Fu, et al. 2016; Zhao, et al. 2017), and altering the steady-state mRNA levels through codon-dependent mRNA decay (Burow, et al. 2018).

Regardless of the particular driving mechanism, the central hypothesis underlying these associations between gene expression and codon usage is that all codons are similarly effective in genes with low expression, and that it is in high expression genes that some are favored and some are disfavored. We examined this hypothesis with our measure of codon fitness *ĝ* by calculating the difference in codon frequency in highly expressed genes and the value in low expression genes and plotting this against *ĝ*. As expected, the codons with the highest positive difference tend to be those with the highest values of *ĝ* (Figure 3A). Figure 3A also shows the previously observed pattern that the GC-ending codons have higher frequencies in high expression genes, as well as higher *g* values.

**Figure 3.**
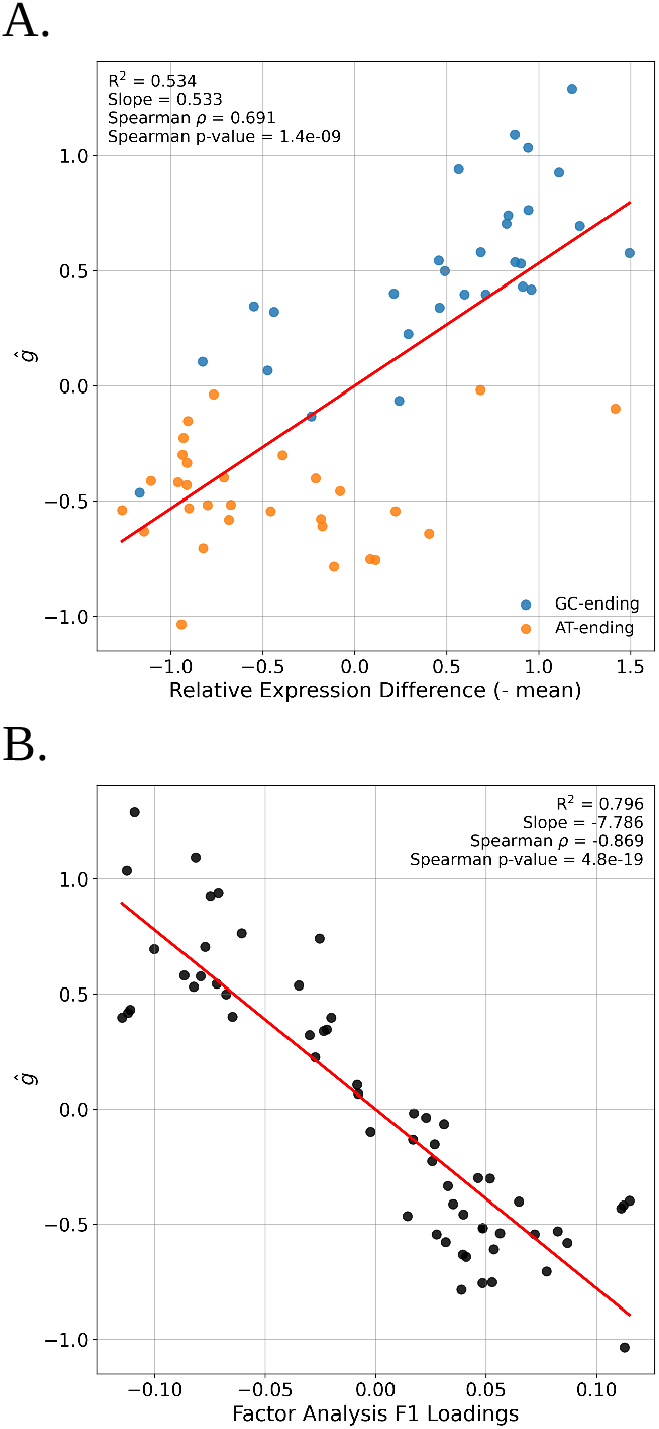
**A.** Correlation between the relative difference in codon abundance in high vs. low expressed genes and codon fitness (*ĝ*). Colors represent codons ending in G/C or A/T. **B**. Correlation between factor analysis’ F1 loadings and *ĝ*.

One way to capture the effect of factors that act similarly on codons across a spectrum of genes is to identify patterns of covariation in codon frequencies across genes. If gene expression is the primary driver, for example, then high expression genes will share a subset of the ‘favored’ codons across the 18 amino acids with multiple codons. This pattern of covariation can be captured directly with a multivariate analysis of codon frequency covariation (Hey and Kliman 2002; Rahman, et al. 2021). We conducted a factor analysis of codon-frequency covariation across the genes of *D. melanogaster* (similar to a principal components analysis and correspondence analysis). The first factor from this analysis reflects the largest axis of covariation in codon frequencies across genes, and if this is driven by natural selection, then we expect to see a strong association between the Factor 1 loadings on individual codons and our measure of codon fitness. Figure 3B shows the association of the first factor from a factor analysis of codon frequency covariation with *ĝ* (the sign of the association, negative in this case, is arbitrary).

Synonymous codon changes can alter transcript half-life if they alter mRNA secondary structure which in-turn affects mRNA stability (Mauger, et al. 2019). At the codon level, it has been shown that different codons are associated with different impacts on mRNA stability, as shown in *D. melanogaster* by Burow et al. (2018). The authors measured decay rates for nearly 8000 mRNAs in *D. melanogaster* embryos and calculated the codon stabilization coefficients (CSC), a measure of the correlation between codon usage and transcript stability (Presnyak, et al. 2015). They found CSC to be positively correlated with a measure of tRNA gene copy number (Burow, et al. 2018). When we examine the association between *ĝ* and CSC, we found a clear and statistically significant positive correlation between CSC and codon fitness (Figure 4A).

**Figure 4.**
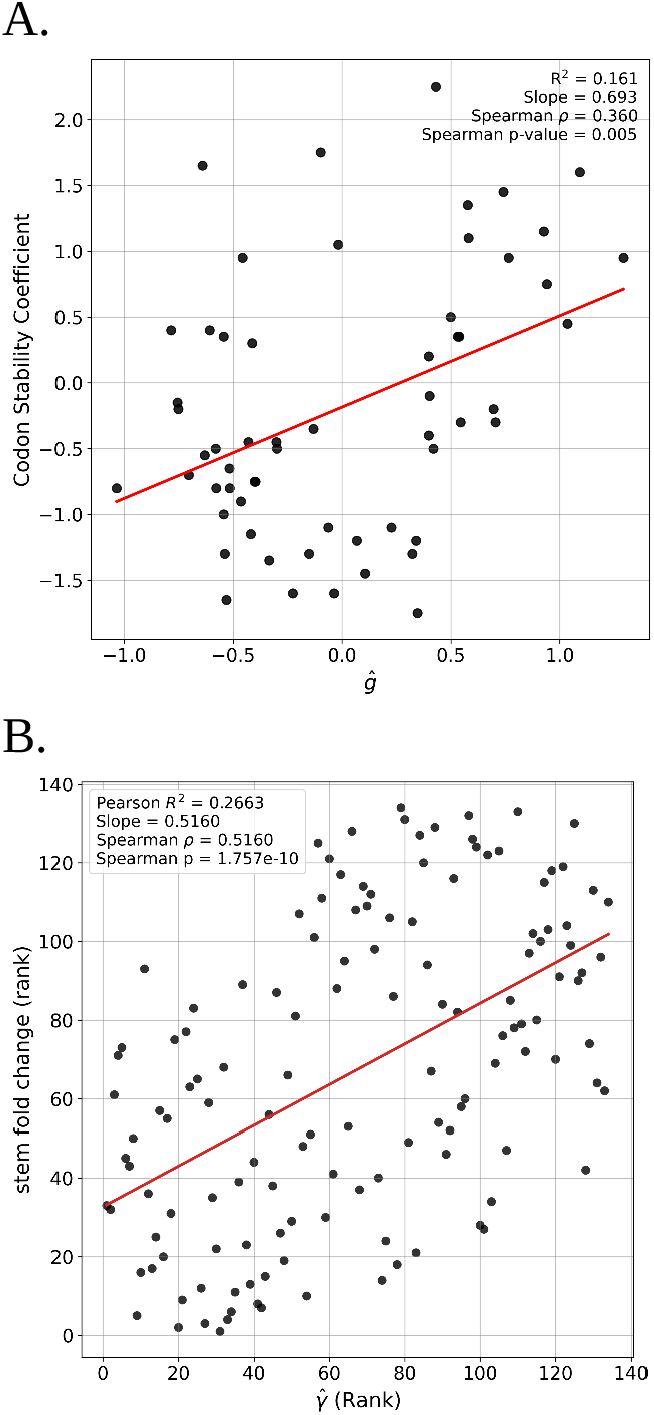
**A.** Correlation of Codon Stability Coefficient and *ĝ*. **B**. Correlation of the rank of fitted selection coefficients and rank of the stem/loop fold change.

We further examined the hypothesis that selection on synonymous codon usage is partly mediated through mRNA folding by estimating secondary structure and recording how often each of the 59 synonymous codons occur in stem and loop sections of the predicted structures. For each of over 12,000 mRNA sequences we estimated secondary structure stem and loop assignments for each base position, and then for each of the 59 synonymous codons determined the degree to which it is overrepresented in stem or loop, relative to mean stem/loop proportions. Then for each synonymous SNP we determined if it occurred in a stem or loop section and if the change to the derived allele was in the direction of a codon that had higher stem representation or loop representation. These values were used to generate a stem enrichment score for each of the 134 ordered codon pairs, which is a measure of the average effect that a specific codon change has on mRNA secondary structure. Model-fitted selection coefficients 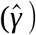 show a highly significant rank correlation with the change in the stem enrichment score of the codon (Figure 4B). The variance explained by this correlation is not high (*R*^2^=0.2663) but is consistent with RNA stability being an important component of codon usage’s impact on gene expression.

## Discussion

Synonymous codon variation has long been thought to be under selection in *D. melanogaster* (Shields, et al. 1988; Sharp and Li 1989; Moriyama and Hartl 1993). However, the strength of this selection has been uncertain, with reports that selection in *D. melanogaster* is undetectable (McVean and Vieira 2001) or that selection is very weak (*γ ⋁* 1) (Akashi 1995), or even quite strong (Machado, et al. 2020).

To better examine the strength of selection we applied a new approach that relies only on ratios of frequencies of segregating sites for candidate selected sites (i.e. synonymous codon changes) and a neutral control (SNPs in short introns) (Hey and Pavinato 2025). For each of 134 pairs of synonymous codons (ancestral/derived) for which the codons are one base apart, we obtained an initial set of estimates of *γ*. We then exploit a key corollary of the hypothesis of weak selection, that a change of synonymous codon A to B will have a selection coefficient that is the additive complement of the change of B to A. When we fit this model to the initial 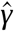 values, we obtained estimates of codon specific fitnesses (i.e., *ĝ*), which we then used in turn to generate a final set of model-fitted 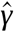 values.

The model-fitted estimates fall in a narrow range 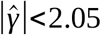. Although the strength of selection is quite modest, it is nevertheless strong enough to shape the site frequency spectra, and to have a large effect on codon frequencies. In particular, the most common codons are GC-ending, and yet these are expected to be at lower frequencies because of the mutational pressure in favor of AT-ending codons. The fitness estimates predicted codon frequencies quite accurately under a simple selection-mutation-drift model, whereas predictions based on mutation rates alone fail to predict codon frequencies. We also observed a strong correlation between estimates of *g* and the first factor from a factor analysis of codon frequency covariation. This observation is consistent with a general model in which gene level selection on codon usage drives patterns of genome-wide variation in codon usage. Values of *ĝ* are also correlated with the difference in codon frequency in highly and lowly expressed genes, as expected if gene expression is a driver of selection on codon usage. Finally, we detect a clear signal that selection on codon usage acts in the direction of stabilizing mRNA secondary structure.

Our estimates do not depend on between species divergence, or on overall codon abundance, or relative codon frequencies, and therefore do not rely on prior designations of “preferred” or “unpreferred”. Moreover, because they are based on ratios of frequencies that are largely insensitive to demographic history—particularly when selection is weak (Hey and Pavinato 2025) —they reflect recent selective forces in the population. Finally the estimates appear robust to ancestral/derived polarization errors based on both the use of higher stringency criteria for ancestral base calls, based on the *D. melanogaster* group whole genome alignment and the use of a deconvolution algorithm to generate simulated data from the actual data under a 5% polarization error rate.

After the initial reports of selection on codon usage in *D. melanogaster* that were based on associations with gene expression (Shields, et al. 1988) and the negative correlation of synonymous substitution rates and codon usage bias (Sharp and Li 1989), there followed a series of reports that found little evidence of widespread selection in *D. melanogaster*. Some of these works found no evidence of selection (McVean and Vieira 2001; Singh, et al. 2007), while others found that selection was very weak, less than estimated here (Akashi 1995, 1996; Nielsen, et al. 2007). Some differences may be a function of statistical power, but the studies reporting no or very weak selection share a reliance on divergence data. Such data are accumulations of processes over much longer spans of time than experienced by polymorphisms, and they therefore include several additional sources of variation. The opportunity for alignment errors is considerably greater in divergence data, and these can greatly distort substitution count or rate estimates (Selberg, et al. 2025). Species divergence data are also affected by changes in which codons are favored by natural selection, as has occurred in *Drosophila* (Bauer DuMont, et al. 2009), as well as by changes in base composition and selection parameters, which have also occurred in the divergence of *D. melanogaster* and related lineages (Akashi, et al. 2006; Singh, et al. 2009). In contrast the approach deployed here, by focusing only on polymorphism frequencies, provides estimates of the forces that have been acting recently in the population. Our results also differ in part from more recent findings that some synonymous codons (e.g. 20%) are under relatively strong selection (e.g. 2 *Ns*>10) (Lawrie, et al. 2013; Machado, et al. 2020). Those findings relied strongly on counts of invariant codon positions, which were not included in this study as invariant sites for specific ancestral/derived codon pairs cannot be identified, and it seems likely that this explains much of the difference in findings.

## Methods

### Polymorphism Data

We extracted synonymous and short intron SNPs from whole-genome sequencing data for the four autosomal chromosome arms of 197 haploid embryo genomes from Zambia (Lack et al. 2015). All sequence data were downloaded from the Drosophila Genome Nexus (DGN, https://www.johnpool.net/genomes.html). We first ran the “masking package” available at DGN to mask identical-by-descent or admixture tracks when present in one or many genomes. Using a custom pipeline that uses the snp-site program (Page, et al. 2016) the multi-alignment FASTA file of the 197 genomes was converted to a VCF file. We used GATK LiftoverVcf to shift the SNP coordinate positions from *D. melanogaster* Dromel 5 (or 3) to Dromel 6 (G dos Santos et al. 2015) and annotated the VCF files with SnpEff (Cingolani, et al. 2012) as done previously (Hey and Pavinato 2025). We rooted the VCF file by estimating the ancestral base using a whole genome alignment of *D. melanogaster* subgroup species (see below). We retained SNPs if they were genotyped in at least 160 genomes and subsampled without replacement SNPs with more than 160 genomes. To examine the effect of using all genomes by imputing the missing data, we used Beagle v.5.5 (Browning and Browning 2007) to build a VCF file with a full sample size of 197. In order to help ensure that the control SNPs were selectively neutral and as closely matched as possible for local mutational context, we focused on short introns (Parsch, et al. 2010) (less than 86 base pairs, with the first and last 8 bases excluded). For each ordered codon pair the intron SFS was assembled by identifying for each synonymous SNP a short intron SNP that had the same reference allele and flanking bases (Machado, et al. 2020; Hey and Pavinato 2025).

### Ancestral Base Estimation

To polarize *D. melanogaster* SNPs by the ancestral base, multiple sequence alignments were generated using Cactus v1.3.0 (Armstrong, et al. 2020). We focused on seven species from the *D. melanogaster* species group: *D. melanogaster, D. simulans, D. sechellia, D. mauritiana, D. yakuba, D. santomea*, and *D. erecta*. Soft-masked reference genome assemblies were obtained from NCBI RefSeq (Supplementary Information Table 1). The guide tree topology and branch lengths were derived from the maximum-likelihood phylogeny of Suvorov et. al. (2022) with *D. santomea* (absent from that study) added as a sister taxon to *D. yakuba* with a branch length of 0.5. The resulting HAL file was converted to multiple alignment format (MAF) file with *D. melanogaster* as the focal species using ‘hal2maf –onlyOrthologs’ to retain only one-to-one orthologous alignments.

Ancestral nucleotide states at polymorphic sites in *D. melanogaster* were inferred using a phylogenetic maximum-likelihood approach. For each biallelic SNP position identified in the Zambia VCF files, we extracted the corresponding alignment block from the MAF file. Only alignment blocks containing sequences from at least five of the seven species were retained for analysis. To avoid circularity in the ancestral state inference, the *D. melanogaster* nucleotide at each SNP position was replaced with a gap character prior to phylogenetic reconstruction. For each alignment block containing one or more SNPs, we ran RAxML-NG v1.0 (Kozlov, et al. 2019) in two stages. First, we inferred the maximum-likelihood tree topology and branch lengths under the General Time Reversible + *Γ* substitution model. Second, we re-ran RAxML-NG with the ‘—ancestral’ flag using the tree inferred from the first run to obtain marginal ancestral state probability distributions at each internal node. We extracted the posterior probabilities for the node immediately ancestral to *D. melanogaster* (i.e., the most recent common ancestor of *D. melanogaster* and its sister clade comprising *D. simulans, D. sechellia*, and *D. mauritiana*). For downstream polarization of SNPs, we required an ancestral state probability of ≥0.9 or ≥0.99 to assign a root; SNPs failing this threshold were excluded from further analyses. This approach yielded ancestral state estimates for approximately 63% of autosomal biallelic SNPs.

### Assembly of selected and neutral SFSs for each ordered codon pair

Site frequency spectra were built for each of the 134 ordered (ancestral to derived) synonymous codon pairs from the polarized SNPs in the Zambia VCF files. Because of some missing data, SFSs were built to a common sample size of 160 by random down-sampling with replacement. For each ordered codon pair SFS a neutral partner SFS of the same total allele count was built by identifying for each synonymous SNP a short intron SNP that had the same reference allele and same flanking bases (Machado, et al. 2020; Hey and Pavinato 2025).

### Simulations of deconvoluted polarization errors

To approximate an error-free site-frequency spectrum (SFS) from data with possible mispolarization, we assumed a uniform mispolarization rate of 0.05 and treated each mirror pair of forward and reverse bins (e.g., TTT→TTC and TTC→TTT; derived-allele counts 1 to 159) as a 2×2 linear mixing system between the forward (*F*) and reverse (*R*)spectra.

Assuming a mispolarization rate (r), the observed forward and reverse spectra are modeled as mixtures of the unseen true spectra. For *N* =160 genomes, *i*=1, ⋯, *N* − 1 and *j*=*N* −*i*: *F*_obs_ [*i*]=(1 −*r*) *F*_true_ [*i*]+*r R*_true_ [*j*] and *R*_obs_ [*j*]=(1 −*r*) *R*_true_ [*j*]+*r F*_true_ [*i*] (and symmetrically for *R*_obs_ [*i*], *F*_obs_ [*j*]. We inverted this 2 × 2 system independently for each mirror pair to obtain deconvolved expectations. The deconvolved forward expectation is *F*_exp_ [*i*]=((1 −*r*) *F*_obs_ [*i*] −*r R*_*obs*_ [*j*])/(1 − 2 *r*)), with analogous expressions for *R*_exp_ [*j*], *R*_exp_], and *F*_exp_ [*j*]. Small negative expectations were truncated to zero. We generated simulated deconvolved spectra by multinomial sampling from these expected values for all 134 pairs of SFSs, while preserving the original number of polymorphic sites in each SFS.

### Fitting a Codon Fitness Model

We consider a model in which selection on the alleles at a synonymous SNP depends only on the codon identities of the segregating alleles, without dominance and without any epistatic interactions with other parts of the genome. Furthermore we suppose that synonymous SNPs are segregating just two alleles (sites with more than two alleles were excluded), in which case the intensity of selection on a SNP can be modelled as the difference in the fitness value of the two segregating codons. Under this model, for an amino acid with two-fold redundancy, the selection coefficient associated with a change in one direction will be the negative of the selection coefficient of the change in the reverse direction. For example, consider all SNPs for the two-fold amino acid tyrosine. When the ancestral state is the codon TAC and the derived state is TAT, we would expect under our simple model that the SFS from all TAC>TAT SNPs could be modelled as being shaped by a selection coefficient *γ*, while the SFS for those SNPs that are TAT>TAC could be modelled using the additive inverse of *γ*, i.e. −*γ*. To examine the prediction that *γ* estimates from both ordered SFSs for a codon pair should be close to additive inverses of each other, forward and reverse pairs were regressed against each other under a model II regression (data associated with both X and Y axes are subject to error) assuming 67 independent points.

For amino acids with *k* >2 codons, there will be *k* × (*k* − 1) ordered pairs of codons and 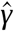 values which, unlike the case of *k* =2, is more than the number of codons. In these cases we can consolidate the multitude of 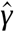 values into a smaller set of codon fitnesses under the assumption that the strength of selection revealed by any particular ordered codon pair is the difference between the ‘to’ codon fitness and the ‘from’ codon fitness. In this model, each codon has a fitness *g* and the strength of selection on mutations from codon *i* to *j* is *γ*_*i, j*_=*g*_*j*_ − *g*_*i*_. As the *g* values include an unknown constant such as variation in mutation rates, … that cancels out when calculating selection coefficients *γ*_*i, j*_, only the relative differences between codon fitnesses matter. For this reason, there are only *k* − 1 *g* values to be estimated, with one codon arbitrarily assigned a reference fitness value.

To fit this model we populate a *k* by *k* matrix with 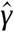 values, set the first codon to have *g*=0, and then estimated the remaining *k* − 1 g values by minimizing the squared deviations between the matrix of 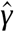 values and the corresponding matrix of *g*_*j*_ − *g*_*i*_ values:

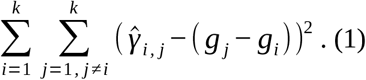

Finally, we subtract from each of the *k* fitness values (including that with *g*=0) the mean 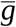 so that the final estimated values have a mean of zero. As the selection intensities depend only on the difference in *g* values, this rescaling retains the basic arithmetic relationship between *ĝ* and 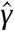 values. Imposing a mean of zero, as assumed under the model, will also act to remove the directional bias observed in the initial 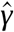 values.

Bootstrap confidence intervals were obtained by sampling pairs of SNPs without replacement to build 120 data sets, each with 134 pairs of synonymous and short intron SFSs. *γ* and *g* estimates were obtained for each simulated data set just as for the actual data, and the 95% confidence intervals estimated from the ordered values for each array of 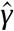 and *ĝ*.

To assess whether the *ĝ* values provide a statistically significant fit to the 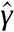 values, for a given amino acid with *k* >2 synonymous codons, we calculated the correlation between the matrix of initial 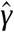 values and the matrix of *ĝ*_*j*_ − *ĝ*_*i*_ values. For amino acids with more than 2 codons, this correlation was compared to 1000 simulated correlations obtained by fitting *g*values for 1000 matrices of randomly shuffled 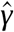 values. The proportion of simulations that provide a higher correlation amounts to a *p* value for the hypothesis that the 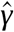 values are associated with each other as specified under the model. For all amino acids with more than two codons the observed correlation was the greater than or equal to either all, or nearly all of 1000 simulations (Supplementary Information Table 7).

### Expected codon frequencies

A number of studies have been conducted to estimate the base pair mutation rates in *D. melanogaster* and if we take the overall mean of these estimates (Assaf, et al. 2017), we can build a matrix of the relative mutation rates between all possible pairs of bases. This can then be used to develop a predicted codon frequencies under an equilibrium model of only mutation, as well as a model with 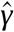 values. Under a Wright-Fisher model with transition rate *q*_*i, j*_from codon *i* to codon *j*, the equilibrium frequency of codon *i, π*_*i*_, is subject to detailed balance such that *π*_*i*_ *q*_*i, j*_=*π*_*j*_ *q*_*j*,*i*_ at equilibrium, which gives the relation

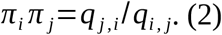

Then given a set of transition rates, and the constraint that 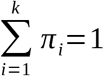, the system of equations of the same form as (1) for all pairs of codons for an amino acid can be reduced to a solution for the equilibrium frequencies.

In a model with weak selection the transition rate from codon *i* to codon *j* is 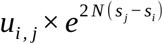 (Ewens 2004) so that 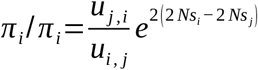. Thus given a set of mutation rate estimates and 2 *Ns* estimates, we can estimate the equilibrium codon frequencies under a selection-mutation-drift equilibrium model. The same framework, with 2 *Ns* values set to a constant can be used to predict equilibrium codon frequencies under a mutation-drift model, i.e., without selection.

### Gene expression data

We obtained all life-stage data for all 12,690 Drosophila genes in the DGET database (Hu, et al. 2017) and summed FPKM (fragments per kilobase million) values across life stages to get an overall gene expression value. Genes were ranked by expression and divided into 104 equally sized bins. Codon frequencies were measured in the lowest and highest bins. For a measure of “difference in codon frequency” we first measured relative difference as the absolute difference in mean frequency among genes in each of the highest and lowest bins, divided by the mean of the frequencies in the two bins. The final measure was this relative difference minus the average of relative differences among the 59 synonymous codons.

### Factor Analysis

Codon frequencies were recorded for each of 59 synonymous codons for 12,905 genes in the *D. melanogaster* reference genome. For each amino acid with multiple codon, these frequencies summed to 1. A conventional factor analysis was conducted of the codon frequencies using the scikit-learn module (Kramer 2016) in python and the first factor (F1) was taken as a measure of codon frequency covariation across genes (Hey and Kliman 2002).

### RNA Secondary Structure Analysis

Secondary structure was estimated for 12,761 mRNA sequences using the ViennaRNA package (Lorenz, et al. 2011).

## Data Availability

SFS data are available in Supplementary Information. Python Scripts are available at https://github.com/vitorpavinato/Dmel-Synonymous-Fitness/tree/main

## Acknowledgments

We are grateful to Michael A. Gilchrist for comments on an earlier version of this paper. This work was supported by National Institutes of Health grant R01GM144468 to JH.

